# Modelling depolarization delay, sodium currents, and electrical potentials in cardiac transverse tubules

**DOI:** 10.1101/611558

**Authors:** Sarah H. Vermij, Hugues Abriel, Jan P. Kucera

## Abstract

T-tubules are invaginations of the lateral membrane of striated muscle cells that provide a large surface for ion channels and signaling proteins, thereby supporting excitation-contraction coupling. T-tubules are often remodeled in heart failure. To better understand the electrical behavior of T-tubules of cardiac cells in health and disease, this study addresses two largely unanswered questions regarding their electrical properties: (1) the delay of T-tubular membrane depolarization and (2) the effects of T-tubular sodium current on T-tubular potentials.

Here, we present an elementary computational model to determine the delay in depolarization of deep T-tubular membrane segments as the narrow T-tubular lumen provides resistance against the extracellular current. We compare healthy tubules to tubules with constrictions and diseased tubules from mouse and human, and conclude that constrictions greatly delay T-tubular depolarization, and diseased T-tubules depolarize faster than healthy ones due to tubule widening. We moreover model the effect of T-tubular sodium current on intraluminal T-tubular potentials. We observe that extracellular potentials become negative during the sodium current transient (up to −50 mV in constricted T-tubules), which feedbacks on sodium channel function (self-attenuation) in a manner resembling ephaptic effects that have been described for intercalated discs where opposing membranes are very close together.

These results show that (1) the excitation-contraction coupling defects seen in diseased cells cannot be explained by T-tubular remodeling alone; and (2) the sodium current may modulate intraluminal potentials. Such extracellular potentials might also affect excitation-contraction coupling.

## INTRODUCTION

Transverse (T-)tubules are deep invaginations of the lateral membrane of skeletal and cardiac muscle cells. In mammalian ventricular cardiomyocytes, T-tubules form a complex network throughout the cell, especially in species with high heart rates such as mice (Pinali et al., 2013; Jayasinghe et al., 2015), and carry many ion channels and regulatory proteins (reviewed in (Bers, 2002; Hong and Shaw, 2017; Bhogal et al., 2018)). Consequently, T-tubules function as a platform for excitation-contraction coupling and signaling, which is essential for the initiation and regulation of muscle contraction (Hong and Shaw, 2017). Importantly, T-tubular remodeling has been reported for several cardiac diseases (Crocini et al., 2017; Crossman et al., 2017a). In particular, T-tubules widen (Wagner et al., 2012; Pinali et al., 2017; Seidel et al., 2017). Understanding the electrical properties of T-tubules in health and disease is therefore paramount to understanding cardiac physiology and pathophysiology. Several questions regarding the electrical properties of T-tubules however remain largely open (Vermij et al., 2019).

A first question concerns the delay after which deep segments of T-tubules depolarize has hardly been assessed. Based on measurements of dextran diffusion out of T-tubules and corresponding modeling of this diffusion process, Uchida and Lopatin recently calculated that T-tubular constrictions and dilations increase the time constant of membrane depolarization from ~10 to ~100 μs, but they did not assess in their experiments the delay of membrane depolarization of deep T-tubular membrane (Uchida and Lopatin, 2018). Since linear cable theory is unsuitable to determine depolarization delay in a structurally complex T-tubule, we present an *in silico* model of a simple T-tubule, an overall constricted tubule, and a tubule with successive constrictions. We quantify the depolarization delay of deep T-tubular segments compared to cell surface, and show that the threshold of voltage-gated channels deep in the cell will be reached slightly later than near the surface.

A second question concerns the role played by T-tubular sodium current. Although the existence of a T-tubular pool of sodium channels is still under debate (Rougier et al., 2019), several studies have suggested that sodium channels are present in T-tubules and that the T-tubular sodium current may be substantial (Maier et al., 2004; Mohler et al., 2004; Brette and Orchard, 2006; Westenbroek et al., 2013; Koleske et al., 2018; Ponce-Balbuena et al., 2018). To date, the effects of tubular sodium current and the interactions between the sodium current and the extracellular potentials have scarcely been investigated. The effects of a large T-tubular sodium current have been already simulated by Hatano *et al.* using an elaborate 3D model of T-tubules without branches, constrictions, and dilations (Hatano et al., 2015). With a sodium current density of 30 mS/μF, the extracellular potential was slightly negative (−1 mV), and sodium current was 8% smaller than at the cell surface. The authors did however not investigate this phenomenon or discuss its physiological importance in further detail. Therefore, in the present work, we extended our model with a T-tubular sodium current. We explore extracellular potentials in our T-tubular model with and without constrictions, investigate the biophysical properties and magnitude of the sodium current throughout the T-tubule, and discuss the physiological implications.

## METHODS

Our model approximated one T-tubule as a cylinder that was divided in 100 segments (for a mouse T-tubule) or 20 segments (for a human T-tubule), separating 101 or 21 nodes, respectively (**Figure 1A, Table 1**). Intracellular resistivity was assumed negligible. T-tubular radius, length, extracellular resistivity, membrane capacitance, and membrane conductance were set as in **Table 1**.

**Figure 1.**
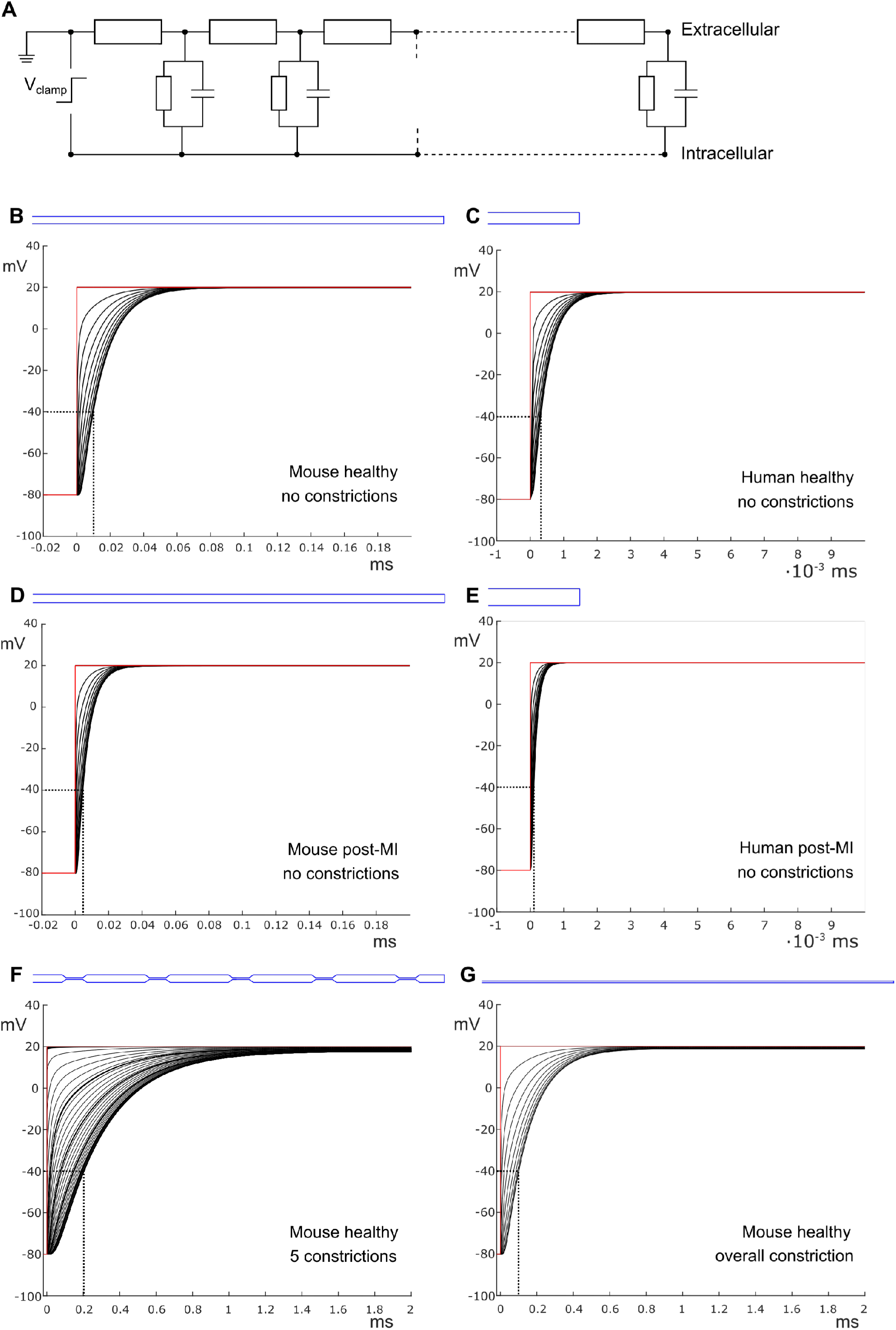
Delay of membrane depolarization in constricted T-tubules. (**A**) Schematic representation of the model. T-tubule is subdivided in approximately 100-nm-long segments consisting of intraluminal resistance, membrane capacitance, and membrane resistance. Cytosolic resistance is assumed negligible and T-tubular mouth is assumed perfectly clamped. Panels **(B-G)** depict simulations of different T-tubules: healthy mouse (**B**), healthy human (**C**), mouse post-myocardial infarction (MI) (**D**), human post-MI (**E**), healthy mouse with five constrictions (**F**), and healthy mouse overall constricted (**G**). T-tubular geometries are schematically depicted in blue and detailed in **Table 1**. (**F**), Curves spaced further apart represent segments of a constriction, curves close together wide segments. Red lines indicate the perfectly clamped mouth of the T-tubule. Note the differences in time scales between mouse with and without constrictions and human. The membrane potentials of every tenth (**B,D,G**) and second (**C,F**) node are depicted. Dotted lines indicate the opening threshold for voltage-gated calcium channels (around −40 mV).

**Table 1.** Parameters for T-tubular model of healthy and post-myocardial (MI) T-tubules of human and mouse. Radius (*r*) and length (*l*) are based on previously published values. Constrictions of healthy mouse T-tubule are also specified: *, parameters of each of the constrictions from the five-constrictions model; **, parameters of the overall constricted model. Numbers between brackets correspond to the following publications: (1) (Wagner et al., 2012), (2) (Hong et al., 2014), (3) (Crossman et al., 2017b), (4) (Cannell et al., 2006). The resistivity of the extracellular space (*ρ*_e_) is set at 100 Ω·cm. Conductance of resting membrane (*g_m_*) is set at 0.143 mS/cm^2^ (Uchida and Lopatin, 2018), which is attributable to a small *I*_K1_ and leak currents. Capacitance (*C*_m_) was set at 2 μF/cm^2^ in healthy cells to simulate microfolds, and at 1 μF/cm^2^ to simulate the loss of microfolds in disease (Page, 1978; Hong et al., 2014).

| Species | $r$<br>(nm) | $l$<br>($\mu\text{m}$) | $\rho_e$<br>($\Omega \cdot \text{cm}$) | $C_m$<br>($\mu\text{F}/\text{cm}^2$) | $g_m$<br>(mS/$\text{cm}^2$) | Nodes |
| --- | --- | --- | --- | --- | --- | --- |
| Healthy mouse | 98.5 <sup>(1)</sup> | 9 <sup>(2)</sup> | 100 | 2 | 0.143 | 101 |
| Constrictions | 9.85 <sup>(1)</sup> | 0.45*/9** | 100 | 2 | 0.143 | 5*/101** |
| Post-MI mouse | 108 <sup>(1)</sup> | 9 <sup>(2)</sup> | 100 | 1 | 0.143 | 101 |
| Healthy human | 147 <sup>(3)</sup> | 2 <sup>(4)</sup> | 100 | 2 | 0.143 | 21 |
| Post-MI human | 218 <sup>(3)</sup> | 2 <sup>(4)</sup> | 100 | 1 | 0.143 | 21 |

In this T-tubular model, sodium current in the different nodes was modeled according to Luo & Rudy (Luo and Rudy, 1991) with modifications by Livshitz & Rudy (Livshitz and Rudy, 2009). A voltage pulse from −85 to +20 mV was applied at the mouth of the tubule (−80 to +20 mV for simulations without sodium current). This situation mimics a cell that is perfectly voltage clamped at the level of its bulk membrane. The value of −20 mV was chosen to elicit a large sodium current. The sodium current maximal conductance was set to 23 mS/μF, the same value as used in whole-cell simulations (Luo and Rudy, 1991). Note that this value is probably on the high side, as the intercalated discs contain a relatively high density of voltage-gated sodium channels and carries a significant proportion of the whole-cell sodium current (Shy et al., 2014). Calcium channels were not included in the model as these channels open only when the majority of sodium channels have already inactivated.

The model was implemented numerically using a one-dimensional finite-difference scheme. Membrane potential (*V*_m_) and the gating variables of the sodium current were integrated using the forward Euler method using a constant time step of 0.25 ns. Simulations were implemented and run in MATLAB (version 2015a, The MathWorks, Natick, MA, USA).

## RESULTS

### DELAY OF T-TUBULAR MEMBRANE DEPOLARIZATION

First, we set out to answer the question how long it takes to charge the membrane as a capacitor and depolarize the T-tubules in the absence of sodium current. In other words: what is the delay between depolarization at the plasma membrane and deep in the T-tubules?

When we consider an electrophysiological experiment in which a cardiomyocyte is voltage-clamped, a voltage step at the pipette site will first induce a capacitive current into the cell membrane, which will cause depolarization. While this current can travel unhindered through the cytoplasm into the T-tubular membrane, the narrow T-tubules will oppose an extracellular “exit resistance” against the capacitive current as it leaves the T-tubule again. The deeper and narrower the T-tubule, the higher the exit resistance, the longer it takes for the T-tubular membrane to depolarize. This delay in membrane depolarization in deep T-tubules can directly affect the ion channels in the T-tubule: the deeper in the T-tubule, the later they can open. According to our model, for a typical mouse T-tubule, the threshold of voltage-gated calcium channels (~ −40 mV) in the innermost node of the T-tubule will be reached after ~10 μs (**Figure 1B**). For a human T-tubule [6], the membrane depolarization would reach the threshold of the innermost calcium channels much faster than for a murine T-tubule, after ~0.3 μs (**Figure 1C**). On a side note, in cardiac disease, T-tubules generally widen [16]. Myocardial infarction (MI) induces an increase in T-tubular diameter of 9% in mice [3], and as much as 33% in humans [5]. Therefore, the “exit resistance” of T-tubules decreases and calcium channels deep in the T-tubule open quicker. For a murine T-tubule after myocardial infarction, the delay to reach the threshold of L-type calcium channels is ~7 μs (**Figure 1D**), and in a T-tubule from human post-myocardial infarction, ~0.1 μs (**Figure 1E**).

For a homogeneous tubule with a cylindrical geometry, classical cable theory (Jack et al., 1975) can be used to predict the voltage drop from the mouth to the end, at steady state, once the capacitive loading is complete. First, one calculates the length constant *λ* as

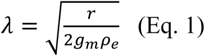

where *r* is the T-tubular radius, *g*_m_ is the conductance of the membrane per unit area, and *ρ*_e_ is the resistivity of extracellular space. It follows that the length constant for the healthy mouse T-tubule (characteristics specified in **Table 1**) is 186 μm. One may then be tempted to use an exponential function (valid for an infinite cable) to describe the decay of membrane potential as

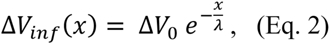

where Δ*V*_0_ = 100 mV is the voltage applied at the tubule mouth and *x* is the distance from the tubule mouth. This approach is however erroneous, because the T-tubule has a sealed end. For sealed-end cables, by applying the reflection and superposition principle, the correct approach is to use a hyperbolic cosine function instead of an exponential (Weidmann, 1952; Jack et al., 1975):

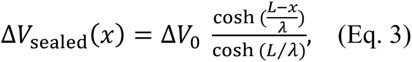

For the healthy mouse T-tubule Eq. 1 yields a voltage drop of 4.72 mV, while Eq. 2 yields only 0.117 mV. This corresponds to the negligible voltage drop depicted in **Figure 1B**. Although the other conditions we modeled lead to different length constants (*λ*_mouse post-MI_ = 194 μm; *λ*_human healthy_ = 227 μm; *λ*_human post-MI_ = 276 μm), the voltage drops are negligible in all cases (mouse post-MI: *ΔV*_inf_ = 0.108 mV, *ΔV*_sealed_ = 0.045 mV; human healthy: *ΔV*_inf_ = 0.877 mV, *ΔV*_sealed_ = 0.0039 mV; human post-MI *ΔV*_inf_ = 0.722 mV, *ΔV*_sealed_ = 0.0026 mV).

### EFFECTS OF T-TUBULAR CONSTRICTIONS ON MEMBRANE DEPOLARIZATION

To quantitatively assess how constrictions change the delay of depolarization deep in a T-tubule, we incorporated constrictions into our model of an unbranched cylindrical T-tubule. We used the parameters for the healthy mouse T-tubule, as these parameters led to the longest depolarization delay (10 μs to depolarize the innermost T-tubule segment to −40 mV). We introduced five 450-nm-long constrictions with a tenfold diameter reduction to 19.7 nm, its centers spaced 1.8 μm apart (**Figure 1F, Table 1**). This is similar to the model of Uchida and Lopatin (Uchida and Lopatin, 2018), which included 20-nm-wide constrictions every 2 μm. For this purpose, the resistances and capacitors depicted in **Figure 1A** were scaled accordingly. With the constrictions in our model, the threshold for Cav channels (−40 mV) in the deepest tubular segment was reached 200 μs later than at the surface (**Figure 1F**). This is 20 times later than in the simpler model without constrictions, where the threshold was reached after 10 μs (**Figure 1B**).

Constricting the tubule ten times to a diameter of 19.7 nm over its full length (**Figure 1G**) increased the depolarization delay in the deep segment 10 times (compare **Figure 1B** and **G**), in agreement with cable theory (capacitive load 10 times smaller and resistance 100 times larger). Interestingly, this increase of the delay was nevertheless smaller than in the presence of five successive constrictions. The difference lies in fact that each widening following each constriction represents a large capacitive load, and the accumulation of these successive loads contributes to slow the spread of electrotonic depolarization.

Additionally, we observed a voltage drop of 2.3 mV in the tubule with five constrictions (**Figure 1F**) and 1.2 mV in the overall constricted tubule (**Figure 1G**), which is ~10 to ~20 times higher than the voltage drop of 0.117 mV calculated for the healthy murine T-tubule.

### IMPLICATIONS OF T-TUBULAR SODIUM CURRENT

As a next step, we investigated the effect of putative voltage-gated sodium (Nav) channels on tubular depolarization (see **Table 1** for parameters). Results using the non-constricted T-tubule, the T-tubule with five constrictions and the T-tubule constricted over its entire length are presented in **Figures 2, 3, and 4**, respectively. **Figure 2B** shows that in a non-constricted T-tubule, the sodium current is smaller deep inside the T-tubule than at the mouth. This smaller current is due to the appearance of a negative extracellular potential in the T-tubule (**Figure 2C**), which results from the flow of current along the narrow tubule. This negative extracellular potential contributes to depolarize the membrane a few mV beyond −20 mV (**Figure 2B**). The extracellular potential becomes a few mV negative at deep T-tubular nodes of the cylindrical mouse tubule (**Figure 2C**), and up to −50 and −40 mV in the tubule with five constrictions and the overall constricted tubule, respectively (**Figures 3 and 4**). This negative extracellular potential brings the transmembrane potential closer to the Nernst potential of sodium (*E*_Na_ = 55 mV). This leads to diminished sodium current (~15% reduction in an unconstructed T-tubule, ~40% in an overall constricted T-tubule, and ~45% in a T-tubule with five constrictions) (**Figures 2D, 3D, 4D**). Thus, the sodium current is smaller in deeper T-tubular membrane segments.

**Figure 2.**
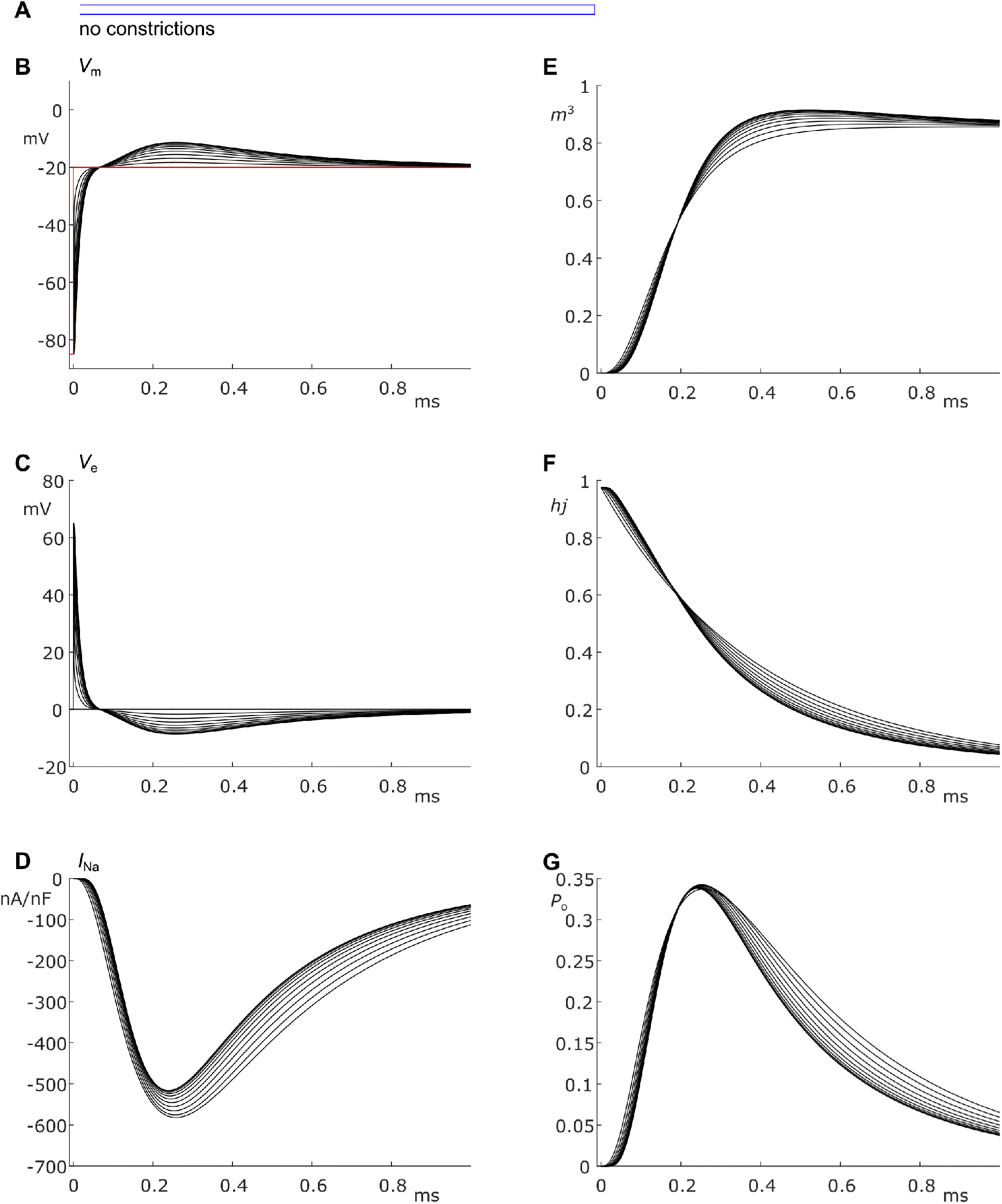
Modeling sodium current in a healthy mouse T-tubule. A voltage-gated sodium current (formulated according to (Luo and Rudy, 1991; Livshitz and Rudy, 2009)) with a conductance of 23 mS/μF (Luo and Rudy, 1991) was introduced to each segment of the healthy mouse T-tubule model in parallel to membrane resistance (see **Figure 1A**). (**A**), Schematic representation of the morphology of a healthy murine T-tubule (see **Table 1**). Membrane potentials (**B**), extracellular potentials (**C**), and simulated sodium current density upon a voltage-clamp step of the tubule mouth from −85 to −20 mV (**D**) are given. Note the decrease of sodium current amplitude and delayed activation in deeper segments of the tubules (**D**). This correlates with changes in the biophysical properties of the sodium current: product of activation gates (*m*^3^, **E**) and inactivation gates (*h_j_*, **F**) show faster activation and inactivation in deeper T-tubular segments, respectively; and peak open probability slightly increases in deeper T-tubular segments (*P*_o_ defined as *m*^3^*h_j_*, **G**).

**Figure 3.**
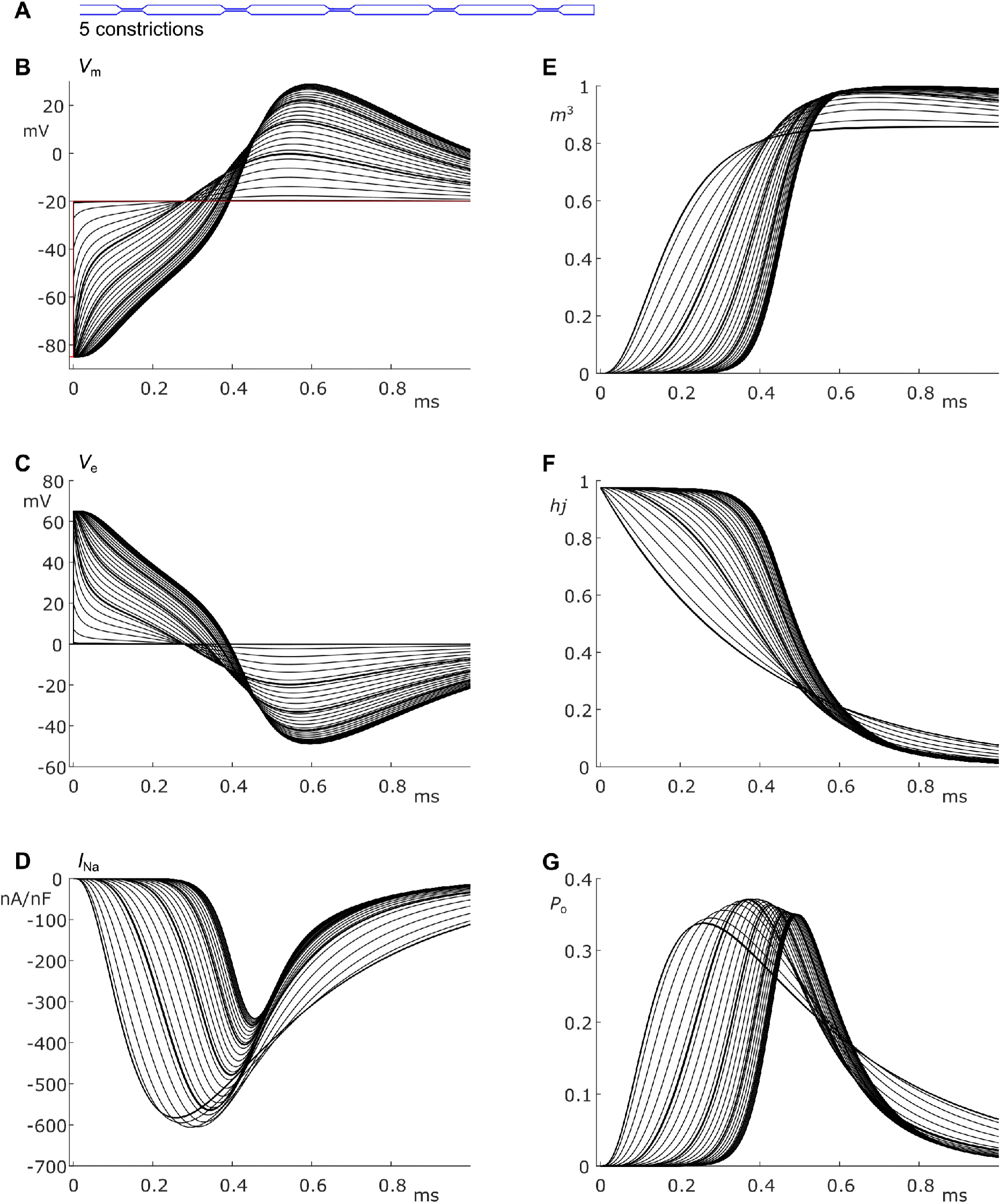
Modeling sodium current in a murine T-tubule with five constrictions. (**A**), Schematic representation of the morphology of the tubule (see **Table 1**). Membrane potentials (**B**), extracellular potential (**C**), and simulated sodium current density upon a voltage-clamp step of the tubule mouth from −85 to −20 mV (**D**) are given. The sodium current amplitude increases over the first constriction and decreases over the other constrictions, while activation is delayed and inactivation is faster in deeper segments of the tubules (**D**). This correlates with a higher driving force (**B**), and changes in the activation gates (*m*^3^, **E**) and inactivation gates (*h_j_*, **F**). Peak open probability slightly increases in deeper T-tubular segments (*P*_o_ defined as *m*^3^*h_j_*, **G**).

**Figure 4.**
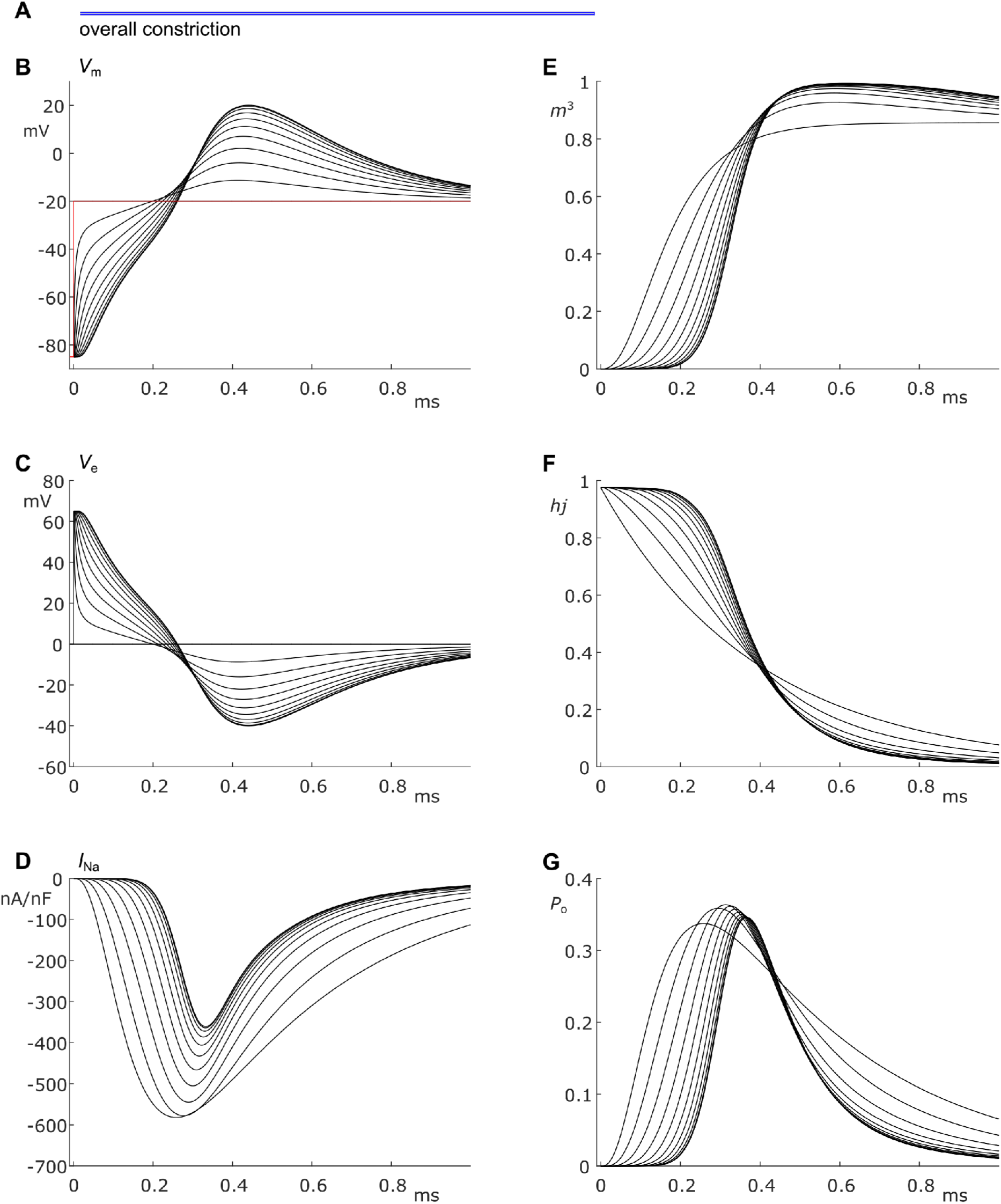
Modeling sodium current in an overall constricted murine T-tubule. (**A**), Schematic representation of the morphology of the tubule (see **Table 1**). Membrane potentials (**B**), extracellular potential (**C**), and simulated sodium current density upon a voltage-clamp step of the tubule mouth from −85 to −20 mV (**D**) are given. The sodium current decreases in deeper segments, while activation is delayed and inactivation is faster in deeper segments of the tubules (**D**). This correlates with a higher driving force (**B**), and changes in the activation gates (*m*^3^, **E**) and inactivation gates (*h_j_*, **F**). Peak open probability slightly increases in deeper T-tubular segments (*P*_o_ defined as *m*^3^*h_j_*, **G**).

Constricting the T-tubule leads to a delay in sodium current activation from 0.03 ms (no constrictions) to 0.15 ms (overall constriction) and 2.7 ms (five constrictions) (**Figures 2D, 3D, 4D**). Interestingly, in the model with five constrictions, the peak sodium current increases over the course of the first constriction (**Figure 3D**). This is explained by the following factors. Firstly, the membrane is not depolarized to −20 mV yet (**Figure 3B**), so the driving force (defined as *V*_m_ - *E*_Na_) is larger. Secondly, the maximal open probability (*P*_o_) in the deep segments is increased compared to the mouth segment (**Figure 3G**), which is a direct result of the voltage-dependence of the activation and inactivation gates (**Figure 3E-F**). For voltage-gated sodium channels, open probability is defined in the Livshitz-Rudy model as *P*_o_ =*m*^3^*h_j_*, where *m* is the activation gate (there are three activation gates), and *h* and *j* the fast and slow inactivation gates, respectively. Since the open probability is only increased by a few percent, the increased driving force contributes more to the increase in peak sodium current.

## DISCUSSION

The results of our computational model show that the delay in T-tubular depolarization and T-tubular sodium current depend on the exact geometry of the T-tubule. The exact location of the constrictions and T-tubular branches are also expected to modulate these factors. We observed that L-type voltage gated calcium channels in deep T-tubular segments attain the activation threshold within a very short time after the cell surface is excited when the “exit resistance” of capacitive current is taken into account (~0.01 ms for a mouse T-tubule without constrictions and up to ~0.2 ms with five constrictions). Thus, the delay of depolarization of mouse and human T-tubules is sufficiently short to ensure excitation-contraction coupling. The T-tubular delay represents no major latency for the sodium current considering that the conduction time along a 100-μm-long cell with a macroscopic conduction velocity of 100 cm/s would be 100 μs (Rohr, 2004).

Human T-tubules depolarize quicker than murine T-tubules because they are relatively wide and short. On a cross-section of a cardiomyocyte, human T-tubules look like spokes on a wheel, and do not form intricate networks like in murine cells (Jayasinghe et al., 2015). The depolarization delay in human T-tubules is therefore negligible in this model. When adapting our model to simulate remodeled T-tubules after myocardial infarction, the activation threshold of calcium channels is reached even faster due to the loss of microfolds (Hong et al., 2014) and increase of luminal diameter (Wagner et al., 2012; Crossman et al., 2017b). The disturbed calcium cycling associated with disease should therefore have other causes, such as dyad uncoupling (Song et al., 2006). The T-tubular widening might however reduce the relative depletion of calcium and accumulation of potassium in the restricted T-tubular lumen, affecting the driving force of their respective ion channels (Hong and Shaw, 2017).

Without dilatations and constrictions, our model gives a negligible voltage drop of 0.117 mV (or 0.117%) at steady state from mouth to deep T-tubular node. This comparable to previously reported results: for a rat T-tubule of 6.84 μm (Soeller and Cannell, 1999), Scardigli *et al.* calculated a length constant of 290 ± 90 μm and a voltage drop from the surface sarcolemma to the core of ~ −4 mV (Scardigli et al., 2017). Scardigli *et al.* however applied the equation for an infinite cable (Eq. 2); applying Eq. 3 for a sealed cable gives a voltage drop of 0.028 mV. A slightly larger but still minuscule value is found for the considerably smaller length constant of 68 μm derived by Uchida *et al.* in a finite element model of dextran diffusion in branched T-tubules with constrictions (Uchida and Lopatin, 2018). When assuming a murine T-tubule of 9 μm, the voltage drop would be 13 mV for an infinite cable and 0.870 mV for a sealed-end cable and is therefore still negligible. However, care must be taken when interpreting these values, since classic linear cable theory cannot be applied straightforwardly to morphologically heterogeneous T-tubules. Indeed, our simulation of a T-tubule with five constrictions show a voltage drop of 2.3 mV, about 20 times higher than the non-constricted tubule. This voltage drop does not affect the opening of voltage-gated channels, which open at much lower potentials.

When inserting the sodium current in our computational model, we found that the sodium current self-attenuates deep in the T-tubules, and the extracellular potential becomes negative (**Figure 2**). This is explained by the very small T-tubule diameter. At the cardiac intercalated disc, where two opposing membranes are also very close together, a similar effect has been predicted (Rhett et al., 2013; Veeraraghavan et al., 2015; Hichri et al., 2018). Importantly, sodium depletion in the T-tubule caused by sodium entering the cell may even augment the self-attenuation of the sodium current because this depletion will decrease the Nernst potential of sodium. Such a phenomenon has been modeled computationally at intercalated discs, but the influence of extracellular potentials nevertheless prevails over sodium depletion (Mori et al., 2008).

The self-attenuation of the sodium current will not affect the calcium current as this effect dies out before the calcium channels open. Interestingly, peak sodium current was increased in the first constriction of our five-constriction model due to a higher channel open probability and a greater driving force. Moreover, sodium current showed faster activation and inactivation kinetics in the deeper segments of the tubule due to the negative extracellular potentials, and the resulting more positive transmembrane potentials.

Given the self-attenuation of the sodium current, it may be interesting to investigate whether the late sodium current in deep T-tubular segments is also quenched, which has also been predicted to occur at the intercalated disc (Greer-Short et al., 2017). However, this process might not play a substantial role in human cardiac arrhythmias, as human ventricular cardiomyocytes contain shorter and wider T-tubules than the murine tubules we modeled here.

Taken together, this study shows that biophysical properties of the sodium current as well as T-tubular depolarization greatly depend on T-tubular geometry. In our model, we did not incorporate elaborate tubular shapes (e.g., tapering of tubules next to constrictions, tubule branching) or ion concentration changes. The first would require a finite element modeling approach and the second would require the implementation of ion fluxes using the Nernst-Planck equation. Both are numerically much more elaborate and demanding. Still, our simple model provides worthy insights into the electrical behavior of T-tubular nanodomains, and we believe that more advanced modeling would yield qualitatively similar results. Should the presence of functional sodium channels be confirmed in the future, it will therefore become of interest to develop such models to obtain a more comprehensive picture.

## AUTHOR CONTRIBUTIONS

Conceptualization (SV), methodology (SV & JK), visualization (SV & JK), writing-original draft preparation (SV), reviewing and editing (SV, HA, JK), supervision (HA & JK), software (JK), formal analysis (JK).

## ACKNOWLEDGMENTS

The authors express their gratitude to Dr. Jean-Sébastien Rougier for thorough feedback on the manuscript.

