## Supplementary material for "Modelling depolarization delay, sodium currents, and electrical potentials in cardiac transverse tubules": Source code: HumanHealthy.m

```
% Simulation of membrane potential in a T-tubule upon stepwise
% depolarization of the tubule mouth

% Vermij, Abriel and Kucera, 2019

% Licence: CC NC-BY-SA

% Human Healthy

clear all
close all

% Parameters
r=0.147 * 0.0001; % radius of tubule (cm)
rho=100; % Resistivity of extracellular space (Ohm*cm)
cm=2; % Capacitance of membrane (uF/cm2)
gm=1/7; % Conductance of membrane (mS/cm2)
L=0.0002; % L: tubule length (cm)

Nnodes=21;

Nsteps=2000000; % Number of time steps
dt=0.00000025; % ms

downs=100; % Downsampling of the output (every "downs"th sample is written to the storage array)

% Voltage clamp parameters at the tubule mouth
vrest=-80;
vstep=20;

%----------------------------

% Compute space constant Lambda

Raxial=rho/(pi*(r^2)); % Ohm*cm/cm2 = Ohm/cm
gmp=gm*(2*pi*r); % mS/cm2 * cm = mS/cm 
rmp=(1/gmp); % kOhm*cm
rmp=1000*rmp; % Ohm*cm
lambda2=rmp/Raxial; % (Ohm*cm) / (Ohm/cm) = cm2
Lambda=sqrt(lambda2); % cm

disp(['Lambda (cm): ' num2str(Lambda)])

%----------------------------

dx=L/(Nnodes-1); % Length of a segment (cm)
Rseg=rho*dx/(pi*(r^2)); % Resistance of a segment (Ohm)
Cseg=2*pi*r*dx*cm; % Capacitance of a segment (uF)

% Configure arrays v_arr (membrane potential) and t_arr (time) to store the results
v_arr=zeros(Nsteps/downs,Nnodes);
t_arr=zeros(1,Nsteps/downs);

% Initial conditions
v=zeros(1,Nnodes); 
v(:)=vrest; % Resting membrane potential (mV)
v(1)=vstep; % Apply vstep at mouth of tubule (first node)
t=0; % time (ms)

%----------------------------
%Loop

for i=0:Nsteps-1
   
    % Write v and t to arrays v_arr and t_arr once every "downs" downsampling steps
    if mod(i,downs)==0        
        v_arr(i/downs+1,:)=v;
        t_arr(i/downs+1)=t;
    end
        
    % Compute currents
    dV=-diff(v); % Potential differences
    iax=dV/Rseg; % Axial current (mV/Ohm = mA)
    diax=diff([0 iax 0]); % difference series of iax; this current must go to the membrane (Kirchhoff's law)
    im=1000*diax/Cseg; % Membrane current (1000 * mA/uF = uA/uF)
    iion=-(v-vrest)*gm/cm; % Ion current density through the membrane; here, assuming a linear current given by gm (mV * mS/cm2 * cm2/uF = uA/uF)
    icap=im-iion; % Capacitive current (uA/uF = mV/ms)
    
    % Integration (Forward Euler)
    v=v-dt*icap;
    t=t+dt;
    
    % Keep the mouth of the tubule clamped at vstep (Dirichlet condition)
    v(1)=vstep;
                                        
end

% Graphical output

figure()
plot(0)
hold on
for i=1:2:Nnodes % every second node
    plot(t_arr,v_arr(:,i),'k')
end
% Plot the voltage clamp step in red
plot([-0.1*max(t_arr) 0],[vrest vrest],'r')
plot([0 0],[vrest vstep],'r')
plot([0 max(t_arr)],[vstep vstep],'r')
hold off
ylim([-100 40])
xlim([-0.001 0.01])
```
