## Supplementary material for "Modelling depolarization delay, sodium currents, and electrical potentials in cardiac transverse tubules": Source code: mhjLR1_livshitz.m

```
function [m_inf,h_inf,j_inf,edtm,edth,edtj,tau_m,tau_h,tau_j] =mhjLR1_livshitz(V,dt)

% This function returns various gating functions for the numerical integration of the 3 gates m, h and j 
% of the Na+ current model of Livshitz and Rudy
% (Biophys J, 2009, 97:1265�1276)

% Inputs:
% V: membrane potential
% dt: time step

% Outputs:
% m_inf,h_inf,j_inf: the steady state values of the gating variables at potential V
% tau_m,tau_h,tau_j: the time constants of the gating variables at potential V
% edtm,edth,edtj: values of exp(-dt/tau) for the 3 gating variables

% Auxiliary "lambda" function as introduced by Livshitz and Rudy
lam=1./(1+exp((V+40)./1.5));

 
% gate m
am = 0.32.*(V+47.13)./(1-exp(-0.1.*(V+47.13)));
bm = 0.08.*exp(-V./11);
m_inf=am./(am+bm);
tau_m=1./(am+bm);
edtm=exp(-dt./tau_m);

% gate h
ah = (1-lam).*(0) + (lam).*(0.135.*exp((80+V)./(-6.8)));
bh = (1-lam).*(1./(0.13.*(1+exp((V+10.66)./(-11.1))))) + (lam).*(3.56.*exp(0.079.*V)+310000.*exp(0.35.*V));
h_inf=ah./(ah+bh);
tau_h=1./(ah+bh);
edth=exp(-dt./tau_h);

%gate j
aj = (1-lam).*(0) + (lam).*((-1.2715*10^5.*exp(.2444.*V)-3.474.*10^(-5).*exp(-.04391.*V)).*(V+37.78)./(1+exp(0.311.*(V+79.23))));
bj = (1-lam).*(0.3.*exp(-2.535e-7.*V)./(1+exp(-.1.*(V+32)))) + (lam).*(0.1212.*exp(-.01052.*V)./(1+exp(-.1378.*(V+40.14))));
j_inf=aj./(aj+bj);
tau_j=1./(aj+bj);
edtj=exp(-dt./tau_j);

end
```
