## Supplementary material for "Modelling depolarization delay, sodium currents, and electrical potentials in cardiac transverse tubules": Source code: MouseHealthyWithINa.m

```
% Simulation of membrane potential in a T-tubule upon stepwise
% depolarization of the tubule mouth
% with Na+ current

% Vermij, Abriel and Kucera, 2019

% Licence: CC NC-BY-SA

% Mouse Healthy with Na+ current (INa)

clear all
close all

% Parameters
r=0.0985 * 0.0001; % radius of tubule (cm)
rho=100; % Resistivity of extracellular space (Ohm*cm)
cm=2; % Capacitance of membrane (uF/cm2)
gm=1/7; % Conductance of membrane (mS/cm2)
L=0.0009; % L: tubule length (cm)

Nnodes=101;

Nsteps=2000000;
dt=0.00000025; % ms

%dt=1; % ms

downs=100; % Downsampling of the output (every "downs"th sample is written to the storage array)

% Nernst potentials of the currents
EK=-85; % for a linear K+ current with conductance gm
ENa=55; % for Na+
gNa=23; % Maximal conductance of the Na+ current (mS/uF)

% Voltage clamp parameters at the tubule mouth
vrest=EK;
vstep=-20;

%----------------------------

% Compute space constant Lambda

Raxial=rho/(pi*(r^2)); % Ohm*cm/cm2 = Ohm/cm
gmp=gm*(2*pi*r); % mS/cm2 * cm = mS/cm 
rmp=(1/gmp); % kOhm*cm
rmp=1000*rmp; % Ohm*cm
lambda2=rmp/Raxial; % (Ohm*cm) / (Ohm/cm) = cm2
Lambda=sqrt(lambda2); % cm

disp(['Lambda (cm): ' num2str(Lambda)])

%----------------------------

dx=L/(Nnodes-1); % Length of a segment (cm)
Rseg=rho*dx/(pi*(r^2)); % Resistance of a segment (Ohm)
Cseg=2*pi*r*dx*cm; % Capacitance of a segment (uF)

% Allocate v (membrane potential), m, h and j (gating variables of INa)
v=zeros(1,Nnodes); % mV
m=zeros(1,Nnodes); 
h=zeros(1,Nnodes);
j=zeros(1,Nnodes);

% Configure arrays v_arr (membrane potential), m_arr, h_arr and j_arr (gating variables of INa) as well as t_arr (time) to store the results
v_arr=zeros(Nsteps/downs,Nnodes);
ina_arr=zeros(Nsteps/downs,Nnodes);
m_arr=zeros(Nsteps/downs,Nnodes);
h_arr=zeros(Nsteps/downs,Nnodes);
j_arr=zeros(Nsteps/downs,Nnodes);
t_arr=zeros(1,Nsteps/downs);

% Setup initial conditions
% v (membrane potential)
v(:)=vrest; % Resting membrane potential (mV)
v(1)=vstep; % Apply vstep at mouth of tubule (first node)
t=0; % time (ms)
% the three gating variables are set to their steady state (inf) conditions at vrest
% mhjLR1_livshitz: function to obtain the gating variables and functions according to Livshitz and Rudy
[m_inf,h_inf,j_inf,~,~,~,~,~,~]=mhjLR1_livshitz(vrest,dt);
m(:)=m_inf;
h(:)=h_inf;
j(:)=j_inf;

%----------------------------
%Loop

for i=0:Nsteps-1
               
    % Compute currents
    dV=-diff(v); % Potential differences
    iax=dV/Rseg; % Axial current (mV/Ohm = mA)
    diax=diff([0 iax 0]); % difference series of iax; this current must go to the membrane (Kirchhoff's law)
    im=1000*diax/Cseg; % Membrane current (1000 * mA/uF = uA/uF)
    iK=(v-EK).*(gm/cm); % K+ current density through the membrane; here, assuming a linear current given by gm (mV * mS/cm2 * cm2/uF = uA/uF)
    iNa=gNa*(m.*m.*m.*h.*j).*(v-ENa); % Na+ current density through the membrane (mV * mS/uF = uA/uF)
    iion=iK+iNa; % Total ion current density
    icap=im+iion; % uA/uF = mV/ms

   % Write v, t, gating variables and iNa to storage arrays every "downs" downsampling steps
    if mod(i,downs)==0        
        v_arr(i/downs+1,:)=v;
        ina_arr(i/downs+1,:)=iNa;
        m_arr(i/downs+1,:)=m;
        h_arr(i/downs+1,:)=h;
        j_arr(i/downs+1,:)=j;
        t_arr(i/downs+1)=t;
    end
    
    % Integration of v (Forward Euler) and t
    v=v-dt*icap;
    t=t+dt;
    
    % Rush-Larsen algorithm for the gating variables
    % mhjLR1_livshitz: function to obtain the gating variables and functions according to Livshitz and Rudy
    [m_inf,h_inf,j_inf,edtm,edth,edtj,tau_m,tau_h,tau_j]=mhjLR1_livshitz(v,dt);
    m=m_inf-edtm.*(m_inf-m);
	h=h_inf-edth.*(h_inf-h);
	j=j_inf-edtj.*(j_inf-j);
    
     % Keep the mouth of the tubule clamped at vstep (Dirichlet condition)
    v(1)=vstep;
                                        
end

% Graphical output

% Transmembrane potential
figure()
plot(0)
hold on
for i=1:10:Nnodes  % every tenth node
    plot(t_arr,v_arr(:,i),'k')
end
plot([-0.1*max(t_arr) 0],[vrest vrest],'r')
plot([0 0],[vrest vstep],'r')
plot([0 max(t_arr)],[vstep vstep],'r')
hold off
xlim([-0.01 max(t_arr)])

% Na+ current density
figure()
hold on
for i=1:10:Nnodes  % every tenth node
    plot(t_arr,ina_arr(:,i),'k')
end
hold off
xlim([-0.01 max(t_arr)])
```
