## Supplementary material for "Modelling depolarization delay, sodium currents, and electrical potentials in cardiac transverse tubules": Source code: README.txt

MouseHealthy.m
Standalone MATLAB script that generates the plot in Figure 1B
(Execution time: approx 20 s)

HumanHealthy.m
Standalone MATLAB script that generates the plot in Figure 1C
(Execution time: approx 20 s)

MouseMI.m
Standalone MATLAB script that generates the plot in Figure 1D
(Execution time: approx 20 s)

HumanMI.m
Standalone MATLAB script that generates the plot in Figure 1E
(Execution time: approx 20 s)

MouseHealthyWithINa.m
MATLAB script that generates the plot in Figures 2B and 2C
This script calls the function mhjLR1_livshitz.m. The corresponding file must be in the same directory.
(Execution time: approx 4 min)

mhjLR1_livshitz.m
MATLAB function to compute various gating functions for the 
numerical integration of the 3 gates m, h and j 
of the Na+ current model of Livshitz and Rudy
Biophys J, 2009, 97:1265�1276
